# Pannotator integrated with Medpipe provides immunological and subcellular location features using a microservice

**DOI:** 10.1101/2024.05.21.595127

**Authors:** Rafael Gonçalves, Anderson Santos

## Abstract

Genome sequencing and assembly are trivial tasks nowadays. After assembling contigs and scaffolds from a genome, the subsequent step is annotation. An annotation evidencing the expected features, like rRNA, tRNA, and CDS, is a signal of the quality of our sequencing and assembly. Different techniques to obtain and reproduce DNA samples, as well as sequencing and assembly of genomes, can impact the quality of a genome’s expected features. The Pannotator tool was conceived as an aid annotation tool focusing on the differences between assembling and its reference genome. Some of the key features for bacterial genome annotations are the subcellular location and immunological potential of a CDS. Instead of reimplementing the prediction of these features in Pannotator, we leveraged the capabilities of our microservice to provide them. In the end, Medpipe software was not modified, and Pannotator underwent minor changes to incorporate the subcellular location and immunological potential of all exported proteins annotated by the tool. Moreover, our Medpipe microservice can also be incorporated into other software. The Medpipe microservice is open to anyone, not only to our Pannotator tool. The successful integration of Medpipe to Pannotator, powered by the Medpipe microservice, offers a powerful approach to advanced genomic analysis. The Medpipe microservice, built on Kotlin with the Spring Boot framework, is instrumental in the automation of Medpipe processing. It achieves this using REST endpoints, such as the execution of Medpipe in an asynchronous manner, status retrieval, and prediction generation, which enhance the modularity and scalability of the microservice. The availability of endpoint documentation, detailed request examples, and logs make our microservice user-friendly. The results of this integration demonstrate the value of the information provided by Medpipe, enriching genomic annotation with additional details, such as the density of mature epitopes (MED) and protein subcellular location classification. The Pannotator has evolved beyond basic function annotation and now provides data on immunological potential, structure, and subcellular location after being integrated with our microservice. The Medpipe microservice is available at https://github.com/santosardr/medpipe-ms.git.

## 1. Introduction

Bioinformatics is an area of research that integrates computer science, statistics, mathematics, and biology. She has been instrumental in advancing research genomics, providing tools and methodologies for data processing and analysis. [1]. In this context, sequence alignment emerged as a fundamental pillar, providing a deep insight into evolutionary relationships and functional disorders between genomes. Biological sequence alignments are tools that, in addition to being used for the analysis of conserved regions and regions that have suffered mutations in homologous sequences, also serve as a starting point for other applications in Computational Biology, such as the study of secondary protein structures and the construction of phylogenetic trees. [2]. The alignment of sequences not only unravels the intricate tapestry of genomic evolution but also serves as a crucial starting point for further investigations. A vital application of this process lies in genomic annotation, where the identification of functional elements, such as genes and regulatory regions, constitutes the essence of the decoding of the genetic code. According to [3], the annotation process is crucial for developing methodologies based on the analysis of genetic material, such as pan-genomics and taxonomies. Genomic annotation is a complex process that requires the identification and cataloging of functional elements in a genome, and the large amount of data generated by sequencing techniques makes this task even more complicated. In this context, the Pannotator (pannotator.facom.ufu.br) is a tool designed to create an automatic annotation from a reference genome. This tool has been developed with the aim of minimizing the workload required in the preparation of and correction of various annotations during the execution of a pan-genome [4]. However, the search for further improvements and predictions has led to the introduction of Mature Epitope Density (Medpipe). Medpipe (medpipe.facom.ufu.br) is a bioinformatics pipeline designed to predict epitope density by mature portions of proteins [5]. Our web pages include Medpipe and Pannotator. In Pannotator, proteins are used to perform the Basic Local Alignment Search Tool (BLAST), which compares proteins from reference genomes. Data on subcellular potential localization (Cytoplasm, Cell Wall, Surface Exposed and Secreted) of proteins generated by Medpipe may be useful to a Pannotator user who is annotating the genome of a bacterium. Until this work, Pannotator users could count on integration with Medpipe to request an annotation service and incorporate it into the annotation of a genome by the Pannotator. Such integration has the potential to reduce the work of a researcher to run the Pannotator and Medpipe separately. In addition, the service offered by Medpipe is unique, and there is no other tool that provides the same service on the Internet. Since Medpipe is a non-free software, open-sourcing is not an alternative. An elegant solution is a communication service between Medpipe and any other biological sequence analysis software. The implementation of a microservice executing and returning the results will meet the expectations of providing additional data for the Pannotator and any other software.

The term “Microservices Architecture” has emerged in recent years to describe a particular way of designing software applications as sets of independently implemented services.

While there is no precise definition of this style of architecture, there are certain common characteristics surrounding the organization of business capability, automated deployment, intelligence in terminals, and decentralized control of languages and data [6]. One of the characteristics of using Microservices Architecture is being able to design a set of independent services, each responsible for a specific part of the functionality, built around business capabilities, which means that they are designed to meet the needs of a particular business. These services deployed independently can be upgraded or retired without affecting the other services. Management is decentralized, which means that everyone is responsible, in addition to being able to write in different programming languages and use different data storage technologies, which gives organizations more flexibility [6].

According to [7], microservices offer benefits such as better scalability, faster deployment, flexibility, resiliency, smaller development teams, modularity, reliability, and reusability. However, there are drawbacks and present challenges in their use, as specified by Velepucha and Flores [8]This new architecture has a high learning curve, requires little experience in team development, has network overhead, is more complex, and duplicates data.

A scalable, high-performance microservice is about more than just the ability to handle a large volume of tasks or concurrent requests. The essence of this lies in its efficiency in performing operations. It can process tasks quickly and effectively, use resources optimally, and maintain high performance even under heavy loads without compromising service quality. It is also prepared for future demand growth by adapting and scaling accordingly. [9]. The flexibility to make modifications to a single service and deploy it in an independently optimized software development and delivery allows the rapid implementation of new features and bug fixes. That autonomy also makes it easier to isolate faults if they occur, restricting the problem to the service in question and simplifying its resolution. In addition, the microservices architecture Enables the reversal of changes quickly and minimizes the risks and negative impacts [10].

Many companies use microservices in their solutions; an example is Netflix, which has developed a microservices platform centered on media workflows, aiming to increase flexibility and speed of development [11]. Another example is Amazon, which faced scalability and productivity issues due to frequent updates, and projects that faced the need to refactor their system and took the microservices approach. The division of monolithic applications into small independent services has enabled the company to respond better to the requirements and make rapid and specific changes [12].

Based on the microservices architecture, we designed the Medµσ that makes use of the Medpipe pipeline. This approach ensures the rapid availability of new features to users. Although the present work shows the integration between Pannotator and Medpipe through the Medpipe microservice (Medµσ), it is essential to point out that the microservice developed is not specific to the use of Pannotator, which can be used by any software or analysis pipeline of biological sequences through the availability of the endpoints on the internet.

The main objective of this work was to create a microservice with the name of Medµσ, which executes Medpipe through an Application Programming Interface (API) following the architecture style Representational State Transfer (REST). We also integrated these APIs with Pannotator, providing a platform accessible to researchers and professionals working with genome assembly and genomic sequence analysis. Specifically, our objectives included describing the development of the Medµσ and integrating the APIs developed in the microservice into Pannotator.

## 2. State-of-the-art Related Research

In this section, we discuss works with objectives like the proposal of the Pannotator and Medpipe.

### 2.1 Review of the Pannotator Literature

The multiplex capability and high throughput of sequencing instruments have made the complete sequencing of the bacterial genome routine. Over the past ten years, several automated genome annotation tools have been made available as open-source software or on publicly accessible pages on the internet. In the following paragraphs, we will concisely address the main features of these tools.

The Prokka is a command-line tool implemented in Perl that allows annotation completion of a preliminary bacterial genome in about 10 minutes on a computer desktop, producing standards-compliant output files for further analysis or visualization in genomic browsers, outperforming web and email-based systems that are unsuitable for sensitive data or integration into computational pipelines [13].

BG7 is an open-source tool based on an annotation paradigm protein-centric gene genomes, designed specifically for bacterial genomes sequenced with next-generation sequencing (NGS) technologies, considering the peculiarities of bacterial genomes (absence of introns and scarcity of sequences non-coding proteins) and NGS technologies, being able to deal with errors and annotate highly fragmented genomes or mixed sequences of several genomes (such as those obtained by metagenomics samples). It has been designed for scalability by using a cloud computing infrastructure based on Amazon Web Services (AWS) [14].

The RAST tool kit (RASTtk) is a modular version of the annotation engine RAST that allows researchers to build custom annotation pipelines, choose software to identify and annotate genomic traits, add features custom to an annotation job, accommodate batch genome submissions, and customize annotation protocols for batch submissions. RASTtk marked the first major software restructuring of RAST since its inception in 2008 [15].

The Protein Sequence Annotation Tool (PSAT) is a meta-platform based on the Web for integrated, high-throughput analysis of genomic sequences, demonstrating its usefulness in annotating the gene products of predicted peptides of *Herbaspirillum sp. strain RV1423*, import the results into the EC2KEGG, and use the resulting functional comparisons to identify a putative catabolic pathway, highlighting the potential in a genome with limited annotation [16]. Genix is a web-based bacterial genome annotation platform, which stands out for providing results closer to the reference annotation, with a lower number of false-positive proteins and non-annotated functional proteins, being able to enhance the accuracy of bacterial genome annotation steps and provide high-quality results [17].

Sma3s is an accurate computational tool for automatic protein annotation with useful functionalities for fundamental and applied science. It provides functional categories and requires low computational resources, allowing complete annotation of proteomes and transcriptomes in about 24 hours on a personal computer [18].

proGenomes is a comprehensive database containing high-quality genomic information from a wide variety of microorganisms. It provides access to bacterial, archaeal, and eukaryotic genomes, along with metagenomes and plasmids. proGenomes is a valuable tool for comparative genomics studies, evolution and microbial research, and new species research [19].

DFAST, a genome annotation pipeline for prokaryotes, also assists in sending data to the public sequence database, with emphasis on its ability to annotate a typical-sized bacterial genome in less than 5 minutes and its integration with the DNA Data Bank of Japan (DDBJ) [20].

EuGene is a gene search tool that can be used in the genomes of prokaryotes and eukaryotes. It uses statistical information, similarities with genes and known proteins, and structured data in GFF3 format to predict the unit transcription of the main genes in the genome and perform functional annotations. This tool can deal with complex genomes with repeating regions and transposable elements and can be configured as ab initio, similarity-based, or hybrid, depending on the sources of information used [21].

DescribePROT is a database that contains 13 descriptors predicted in amino acid level for protein structure and function, encompassing 83 proteomes complete model organisms and including 7.8 billion predictions for 600 million amino acids in 1.4 million proteins, with the possibility of searching for amino acid sequences and UniProt accession numbers [22]. μProteInS is a proteogenomics pipeline implemented in Python 3.8. It combines genomics, transcriptomics, and proteomics to identify microproteins in bacteria, overcoming the limitations of traditional approaches and enabling the identification of Small ORFs (smORFs) with overlapping genes, leaderless transcripts, and sequences not preserved. μProteInS is distributed as open-source software [23].

### 2.2 Review of the Literature on Medpipe

We present here a review of the literature on software and web servers that have been available for the last ten years to find candidate proteins for vaccines or diagnostic targets against pathogenic bacteria. Surprisingly, studies have yet to be conducted on this theme.

Jenner-Predict is a web server that uses a knowledge-based approach to bacterial pathogenesis to predict protein-based vaccine candidates (PVCs) from proteomes of bacterial pathogens. The web server considers domains of different classes of proteins involved in host-pathogen interactions and pathogenesis, including adhesins, virulence, invasins, porins, flagellin, toxins, and others. In addition, Jenner-Predict evaluates the potential immunogenicity of PVCs, comparing them to known epitopes and considering the absence of autoimmunity and conservation in different strains. The server has demonstrated high accuracy in predicting known PVCs and overcame existing methods such as NERVE, Vaxign, and VaxiJen [24].

Another site, VacTarBac, is a web server that uses an immunoinformatic approach to identify vaccine candidates based on epitopes against 14 species of pathogenic bacteria. The server utilizes a comprehensive analysis of target proteins, including virulence factors and essential genes, to predict epitopes with the potential to stimulate different components of the immune system. In addition, VacTarBac removed self-recognized epitopes to prevent unwanted immune responses. The analysis revealed 21 proteins from 5 bacterial species as targets of promising vaccines. The server also identifies B-cell epitopes, T cells, and MHC-II ligands, as well as adjuvants, resulting in a total of 252 unique epitopes. VacTarBac features visualization to assist users in identifying the best vaccine candidates in an antigenic sequence [25].

The Integrative Vaccine Investigation and Online Information Network (VIOLIN) is a database and vaccine research analysis system that curates, stores, analyzes, and integrates diverse vaccine-related research data. Since its first publication in 2008, VIOLIN has undergone significant updates. It currently includes more than 3240 vaccines for 192 infectious diseases and eight non-infectious diseases. Within VIOLIN, there are more than ten independent programs. As an example of programs, we can cite Teeth, such as Protegen, which stores antigenic proteins that have been proven to be valid for vaccine development, and VirmugenDB, which annotates virulence factor genes that can be mutated to generate attenuated vaccines successfully. The VIOLIN also includes Vaxign, the first vaccine candidate prediction program based on reverse vaccinology, and other components of vaccines, such as adjuvants (Vaxjo) and DNA vaccine plasmids (DNAVaxDB). In addition, VIOLIN has databases of licensed human vaccines (Huvax) and veterinary vaccines (Vevax). The Ontology of Vaccines is applied to standardize and integrate the different data in VIOLIN. The VIOLIN also hosts the Vaccine Adverse Event Ontology (OVAE), which represents adverse events associated with licensed human vaccines [26].

A non-web-based alternative is TiD, a stand-alone application that identifies potential targets for drug development. It uses the premise that a protein must be essential for the survival of the pathogen and not homologous to the host to qualify as a target. TiD removes paralogous proteins, selects essential organisms, and excludes those that are homologous to host organisms. Targets are classified as known, new, or virulent. Users can perform road analysis metabolic, protein interactions, and other functionalities through integrated web servers. Targets identified by TiD for Listeria monocytogenes, Bacillus anthracis, and Pseudomonas aeruginosa showed overlap with previous studies. TiD is a useful tool for the rational development of medicines, as it analyzes targets in a bacterial proteome in about two hours [26].

### 2.3 Annotation and Pathogenicity Integration

In addition to providing functional annotation, some tools are also available to provide additional annotation data regarding the type of protein export, resistance to antibiotics, structure, and immune capacity of proteins. For example, MacSyFinder is a bioinformatics tool that allows the search for genetic systems in whole genomes. It utilizes a standards-based approach and Domain profiles to identify and annotate secretion systems, antibiotic resistance systems, and other genetic systems in different organisms. The MacSyFinder is distributed as an open-source software [27].

In the chapter “Antigen Discovery in Bacterial Panproteomes,” an in-silico methodology is described which integrates pan genomic, immunoinformatic, structural, and evolutionary approaches to screening for potential antigens in each bacterial species, aiming at the development of broadly protective vaccines and avoiding specific immunity to alleles, in addition to allowing the development of diagnostic assays [28].

The purpose of our work was to follow these software examples, integrating the Pannotator with Medpipe and providing, along with functional annotation, data that helps to find proteins in a genome with potential for vaccine production and testing of pathogen diagnostics. However, our proposal went further when it implemented a protein pathogenicity annotation microservice that any other genomic annotation software can use. Our software, rather than a competitor, is a functional annotation software, which proposes to be a partner of the most functional annotation capabilities that currently exist when offering a microservice that can be consumed democratically.

## 3. Methods

In this section, we will detail how the Medµσ was conceived, designed, and built, as well as the technologies used and the implementation of the crucial endpoints.

### 3.1 Microservices Architecture

Microservices are built around businesses and can be deployed independently [6]. As shown in Figure 1, each microservice can be responsible for a specific biological analysis or query and function independently without affecting the others. In this example, MS-1 could be a service for running genetic analysis tools on a DNA sequence, MS-2 is a service that performs protein structure prediction using a protein sequence, and MS-3 is a service for searching for specific gene sequences in a genome. The three microservices are independent, so it makes it easy to make any changes to any of them. If the need arises to add a new genetic analysis tool, only MS-1 will be affected without any side effects from MS-2 and MS-3. As shown in Figure 1, each microservice can be responsible for a specific business and function independently without affecting the others. In this example, MS-1 could be a registration service for employees of a company, MS-2 is a service that performs tax calculations for the company, and MS-3 is a service for supplier registration of the company. All three microservices are independent, so it makes it easy to make any changes to any of them. If the company needs to add a new tax calculation rule, only MS-2 will be affected without any side effects from MS-1 and MS-3. In the case of Medµσ, which was built for this work, we followed the same logic. If it is necessary to run or search for results from a bioinformatics pipeline other than Medpipe, another microservice should be built. The Medµσ was constructed only for the execution and search of Medpipe results. Any new functionality that is not part of Medpipe should not be included in the Medµσ, from which one should think about the construction of a new microservice.

**Figure 1.**
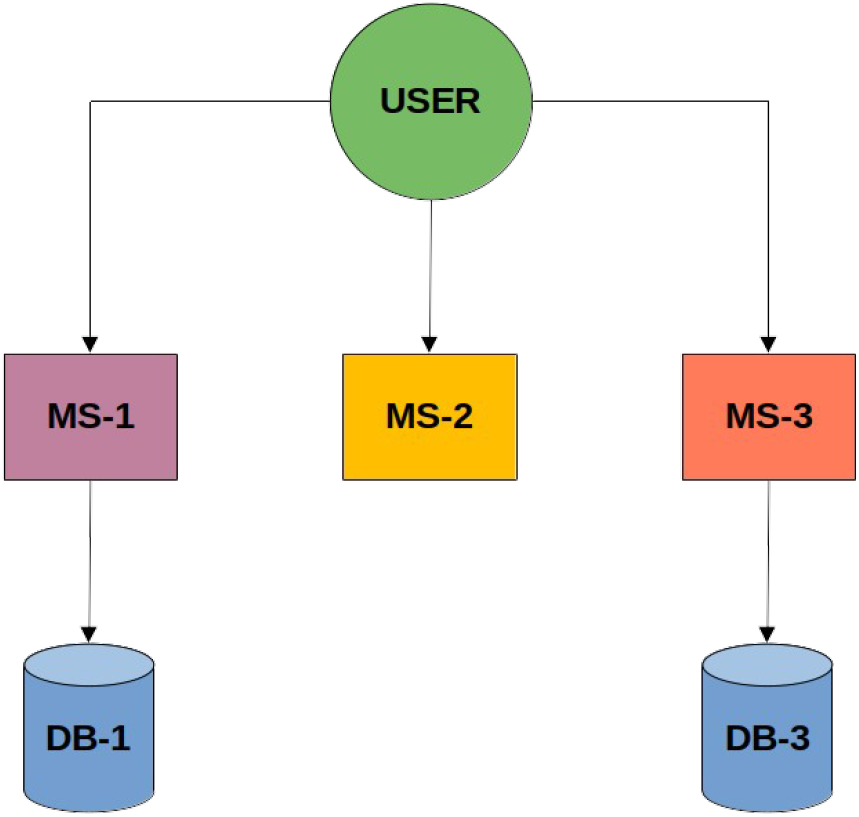
Example of microservices architecture.

We followed the same logic in the case of Medµσ, which was built for this work. Medµσ was constructed only for the execution and search of Medpipe results. Any new functionality that is not part of Medpipe should not be included in Medµσ, from which one should think about the construction of a new microservice. Figure 2 depicts the Medµσ schema.

**Figure 2.**
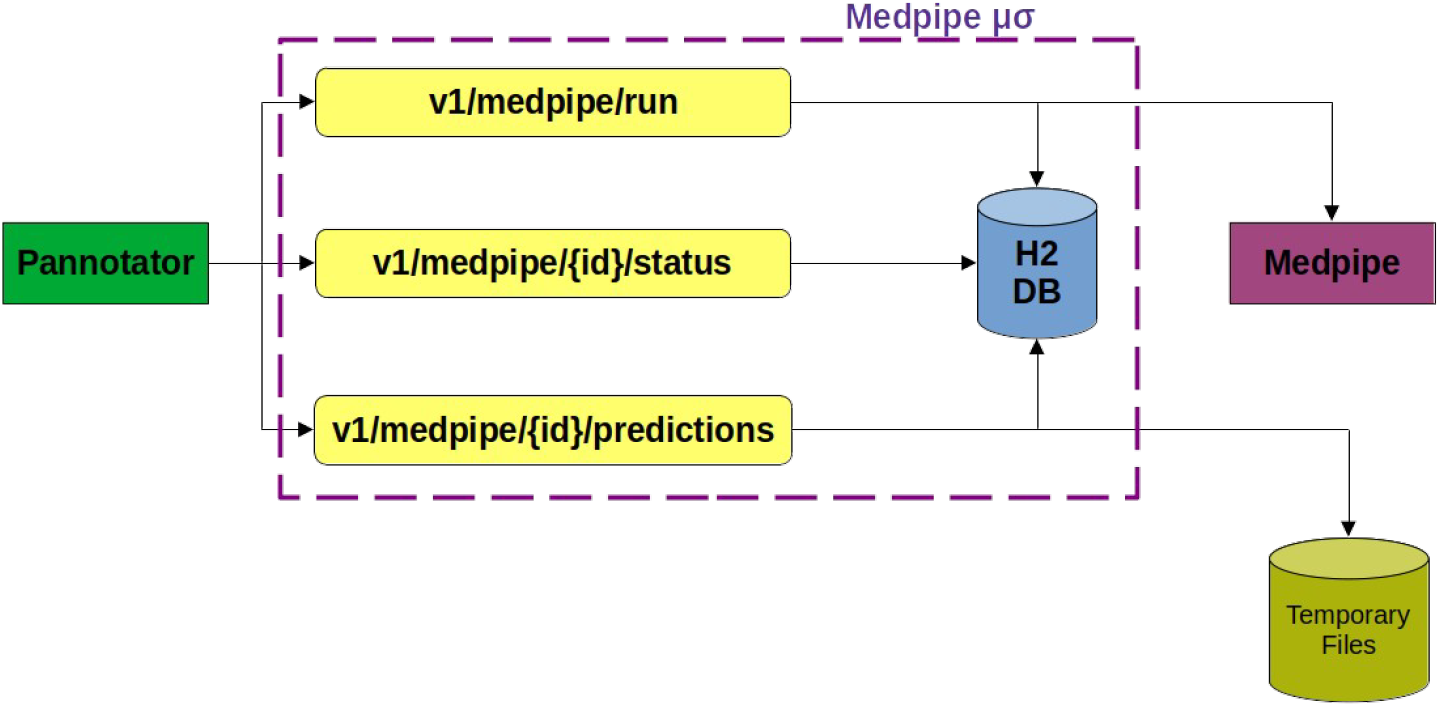
Medpipe microservice architecture. The Pannotator integrates with the Medpipe microservice using the endpoints. Pannotator and Medpipe are basically bash scripts that can be run from the command line or through their web interfaces. With the development of our microservice, Pannotator now runs Medpipe through an API call that is exposed on the internet. The execution of Medpipe is done by the /v1/medpipe/run route and has been included in the Pannotator after updating the target fasta file.

The supplementary material documenting and instructing how to use Medµσ in your bioinformatic tool is accessible in the README file from the GitHub repository at https://github.com/santosardr/medpipe-ms.git.

### 3.2 Programming Language

Kotlin was chosen for microservice development. Created by JetBrains, Kotlin is a modern programming language that is secure, expressive, and interoperable with Java [29]. Compared to Java, Kotlin minimizes the need for boilerplate code, resulting in clearer, more maintainable code.

### 3.3 Spring Boot

Spring Boot is a Java application development framework that is based on the principle of “only what is necessary.” It provides a few features and tools needed to build Java applications, but it only offers essential features. Right makes Spring Boot a lightweight and easy-to-use platform [30]. One of Spring Boot’s outstanding features is its “opinion on configuration” approach. This approach provides sensible default settings for many application aspects, allowing developers to focus more on business logic than on complex configurations. In addition, Spring Boot offers an integrated system for building and managing dependencies, which simplifies the management of the project’s required libraries.

### 3.4 Maven

Maven is a built-in automation and project management tool based on an artifact model. An artifact model is a framework that defines the artifacts a project can have, such as source code, libraries, configuration files, and other files. Maven uses the Artifact template to automate the build, test, and packaging processes and project deployment [31].

### 3.5 H2 Database

The H2 database, a relational database written in Java, was chosen to store information related to the integration processes. It can be Run in client-server mode or embedded mode. In client-server mode, the database runs on a separate server from the Java application. In inline mode, the database runs in the same process as the Java application [32].

## 4. Results

Medµσ provides detailed information on mature epitope density and the protein’s classification in relation to its component in Gene Ontology. To depict the annotation improvements provided by Medµσ in Pannotator, we selected a region from one of our in-house genomes annotated using Pannotator. The genome region was chosen because it illustrates several situations where one can visually perceive the advantages of our visualization schema proportioned by our tool Pannotator and the addends amended by the Medµσ. The illustration in Figures 4 and 5 encompasses several membrane integral proteins, one protein potentially exposed at the bacterial membrane surface, another pool of cytoplasmic proteins, and a single secreted protein. Moreover, three proteins possess low amino acid identity compared to the proteins of the reference genome to the species, eleven proteins with more than 95% amino acid identity and size compared to the same proteins in the reference genome, and two standing between 70 and 94% of identity and size compared to the orthologues present in the reference genome. However, all this information is absent in Figure 3, the standard genome representation for GenBank files.

**Figure 3.**
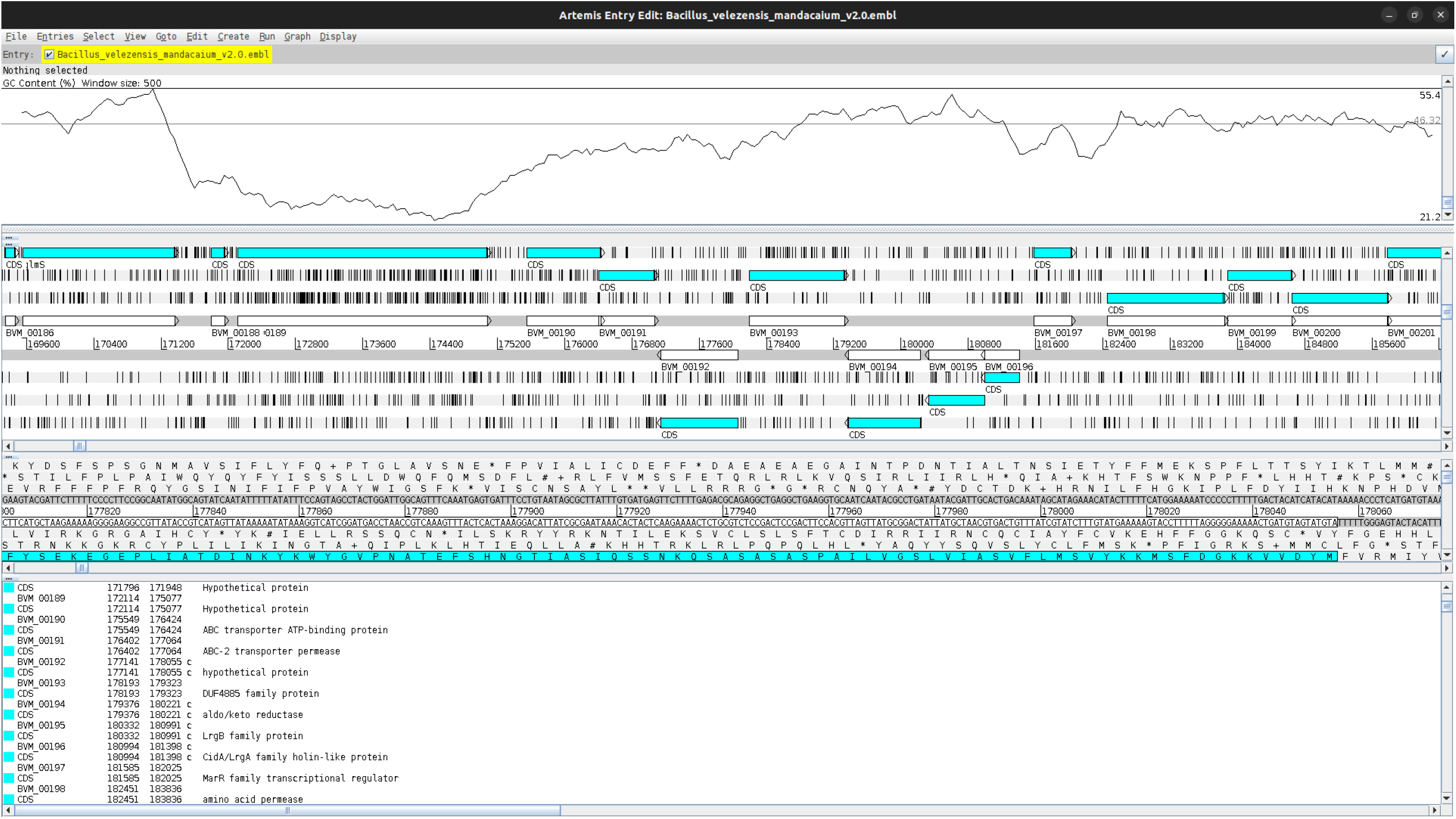
Result Only with Pannotator annotations. The standard genome representation lacks lots of feature annotations from users, making it difficult

**Figure 4.**
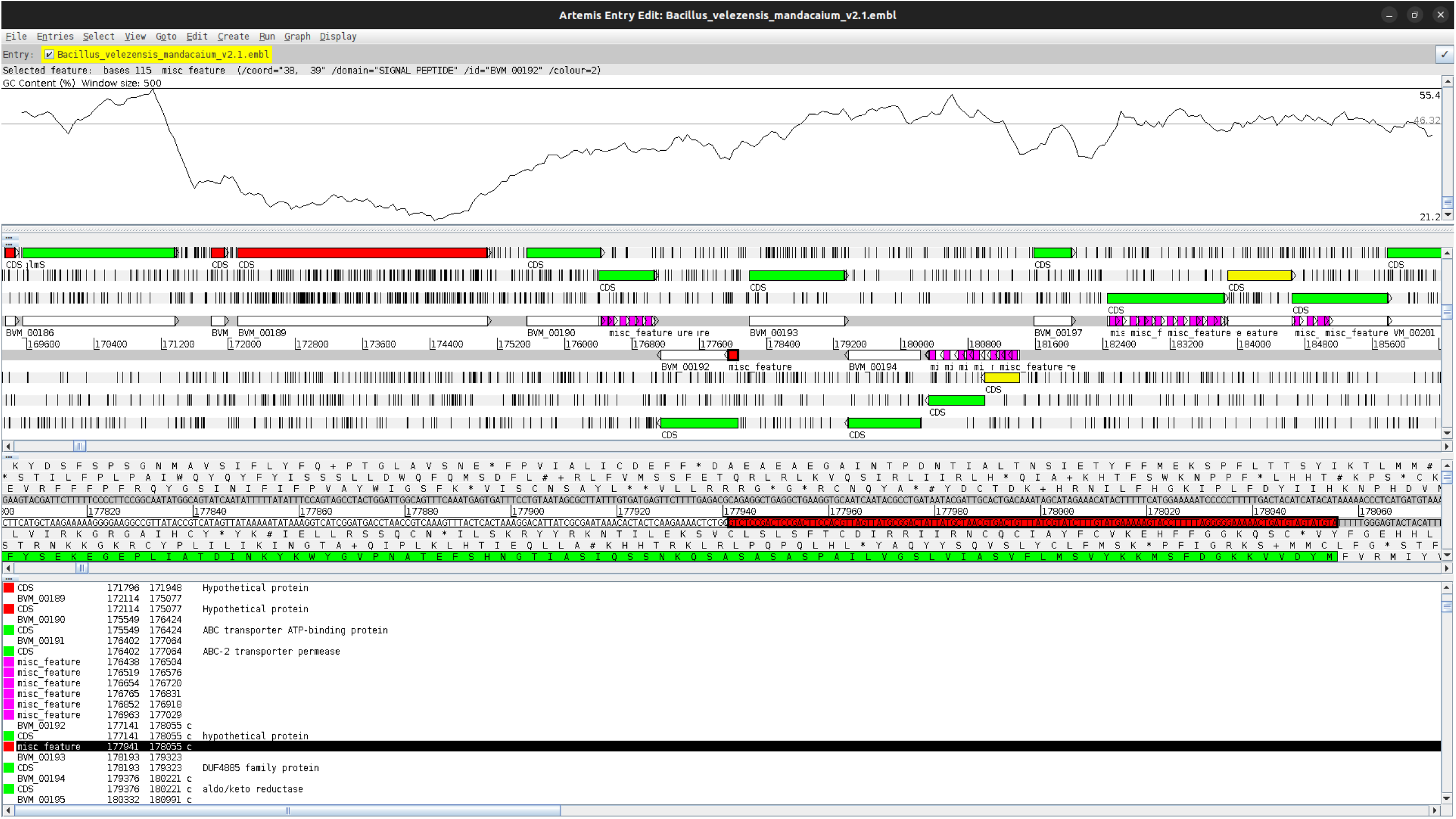
Medµσ annotations included in the Pannotator. The Pannotator color code: green (highly identical to a protein from the reference genome), yellow (most likely to the reference genome), and red (most unlikely to reference genome). The red ones correspond to the below-mean GC content. Signal peptides and transmembrane domains are represented in the DNA strands corresponding to the proteins in red and magenta, respectively.

The only information we provided here that one could infer by analyzing Figure 3 concerns two genes with a below-average GC content region (the third and fourth genes in the forward strand). These two genes can be considered atypical to the reference genome, and one can infer low protein identity to the reference’s proteins.

The features we informed are coded as a color scheme in Figure 4. The coding sequences possessing more than 95% identity and size compared to their best Blast matches in the reference genome are green-colored, the yellow-colored proteins are between 70 and 94% identity and size to the references’ proteins, and the red-colored proteins are below 70%. The 70% is the default cut-off value adopted by our Pannotator tool to apply this color code to all annotated proteins. We should note that our default value of 70% is appropriated to unmask genes possessing unconventional GC content since the genes colored in red fall within the region of below-average GC content. This coloring schema is used by the Pannotator tool as a default. If a user does not want to keep this color code (for NCBI submission purposes), a simple color tag removal drops the color code from a genome annotated by Pannotator.

Besides an efficient color code for coding sequences, the Pannotator can now incorporate the local subcellar predicted to all coding sequences by representing hydrophobic regions indicated by the Medµσ. The Pannotator stores this data projection in the DNA strand corresponding to the coding sequence, using the GenBank features called ‘misc_feature’ or miscellaneous feature. Hydrophobic regions representing signal peptides, a secretion signal, are red-colored. However, this color does not conflict with red-coding sequences since one data is written in the DNA strand and the other in the reading frames. The hydrophobic regions occurring in batches through DNA strands are a signal of membrane or surface-exposed proteins. One can differentiate both according to the number and location of the hydrophobic regions. For instance, in Figure 4, the six coding sequences located at the forward strand have hydrophobic domains along all its extensions, a signal of membrane integral proteins. However, the second protein from the right to the left has no hydrophobic regions covering all the protein extensions, allowing for the possibility of a surface-exposed protein. The remainder of proteins in Figure 4 can be cytoplasmic since no hydrophobic domains have been predicted along the entire protein extensions.

Other data incorporated into the Pannotator results by Medµσ is a statistic depicting proteins more prone to success in crafting a vaccine or diagnostic test for pathogenic bacteria. This statistic is called Mature Epitope Density (MED). An article by the principal investigator of this work first defined the MED [5]. The idea is that secreted and surface-exposed proteins are more prone to strongly containing binder epitopes to the MHC molecules. As many epitopes are predicted in a protein, the odds of starting a host’s immunological response to infections are greater. The coding sequences annotated by the Pannotator and predicted as secreted or surface exposed by Medpipe receive a GenBank feature note indicating the MED statistic. This data is normalized by the greater MED value obtained among all secreted and surface-exposed proteins, so the better targets for a vaccine have the MED close to the value of one. To consult the MED using Artemis, users can ask the software to show the properties of coding sequences containing a few hydrophobic motifs (transmembrane and signal peptide motifs) at the beginning of a coding sequence. The Artemis user will find in the note feature a text like this: “Mature Epitope Density (MED): 1.0” or other values. To avoid a blind search strategy, an Artemis user can also ask the software to list all proteins contained inside the note feature with the keyword “MED.” One should pay attention to the previous description of “a few hydrophobic motifs” that we used in the last sentence. The reason is that a coding sequencing with many such motifs is probably not secreted or surface-exposed but is membrane integral. All membrane proteins have larger MED statistics, even greater than secreted and surface-exposed proteins. However, if we are trying to produce a vaccine using membrane proteins, this task promises to be harder, considering wet lab techniques to isolate and express these proteins. Because of the enormous obstacles to using membrane proteins as vaccine targets, we opt not to show MED for membrane proteins. We also choose not to include cytoplasmatic proteins due to low MED and being most of a predicted proteome, which poses extra and unjustified load to our hardware executing the MED predictions.

## 5. Discussion

The Medpipe integration with Pannotator enriched our genomic analysis, providing additional information that is essential for understanding the biology and functionality of the organisms under study. This data has the potential to drive future scientific discoveries and research, contributing to a more comprehensive and detailed understanding of genomics and proteins of interest.

The fact that the literature review on Medpipe-like tools in the last ten years has returned so few results makes us endorse the hypothesis that pharmaceutical companies would not be interested in implementing vaccines against many of the infectious diseases transmitted by bacteria for which we have today’s antibiotics. The development of a vaccine can take decades and involve billions of dollars in spending (RAPPUOLI et al., 2018). The reason for this lack of interest in solutions against infectious diseases would be in the sale of antibiotics and anti-inflammatories, one of the main pillars of support for the pharmaceutical industries, and the employability of physicians who, in theory, control the release of these drugs. The sale of antibiotics never stops, while a few doses of a vaccine mean the end of the trade of millions or billions in antibiotics. Another probable reason is the fact that many bacterial diseases impact underdeveloped or developing countries (diseases neglected by rich countries). The main stakeholders in vaccines are countries that cannot create them. Still, these countries will be long-standing customers of pharmaceutical companies by buying antibiotics and anti-inflammatories.

## 6. Conclusion

The integration of Medpipe into Pannotator represents an advancement in the ability to analyze and interpret genomic data. The results obtained show the effectiveness of this integration by providing detailed data on gene annotation and genetic products, including Mature Epitope Density (MED) and classification of the protein in relation to its component in Gene Ontology (GO).

The automation of the process, provided by the integration with the microservice Medµσ, simplifies and streamlines genomic analyses. The applicability of the integration extends to projects and research that require a deeper, more comprehensive understanding of genomic data. The availability of Medµσ endpoints capabilities allows for future expansions and integrations with other tools and services, providing a flexible and adaptable platform to the needs of ever-evolving genomic research. Therefore, the successful integration of Medpipe into Pannotator using a microservice called Medµσ represents an advancement in capacity analysis and interpretation of genomic data, providing additional data to a more efficient genomic annotation.

## Supporting information

README

## Supplementary data

It is not applicable to the scope of this article.

## Acknowledgment

The authors confirm that the supplementary data is available in this article.

## Conflicts of interests

The authors declare that there are no conflicts of interest regarding the publication of this article.

## Ethical statement

It is not applicable to the scope of this article.

## Authors’ contribution

Conceptualization, A.S.; methodology, A.S., and R.G.; software, A.S., and R.G.; validation, A.S.; formal analysis, A.S.; investigation, A.S.; resources, A.S.; data curation, A.S.; writing —original draft preparation, A.S.; writing—review and editing, A.S.; visualization, A.S., and R.G.; supervision, A.S.; project administration, A.S. All authors have read and agreed to the published version of the manuscript.

## Notes

https://github.com/santosardr/medpipe-ms.git

### Competing Interest Statement

The authors have declared no competing interest.

