## Supplementary material for "Pannotator integrated with Medpipe provides immunological and subcellular location features using a microservice": README

This microservice was developed using the Kotlin language to deliver the local subcellular and the Mature Epitope Density predicted by the software MEDPIPE for a protein multifasta file. This software is a partial requisite for Rafael Gonçalves' final paper at the Federal University of Uberlândia, Brazil

### Supplementary Material

#### Running the Medpipe microservice

A good thing about microservices is that you can have multiple servers running them. In case of down-service of one, we can try another. I intend to have at least three servers providing the Medpipe microservice. To change to a new server, you should alter the "server" variable in the script. The below script sends the target file to the server. After finishing the processing, three reports will be returned and printed.

```
#!/bin/bash
server="bioinfo.facom.ufu.br"
email="" #This the sender address. You can try your own e-mail to hav

dirfastaFile=target.fasta # Do not alter this file name
cellWall="65" # Measure in amino acids
organismGroup="1" # 0=gram-negative, and 1=gram-positive bacteria
medpipePostURL="curl --location "$server"/v1/medpipe/run --form file=@$dirfastaFile --form c
echo "URL: $medpipePostURL"
processId=`$medpipePostURL`
echo "Result: $processId"

getStatusUrl="curl --location $server/v1/medpipe/$processId/status"
statusexec=`$getStatusUrl`
echo "status: $statusexec"

while [ $statusexec -gt 0 ]; do
    echo "Status: $statusexec"
    sleep 20
    getStatusUrl="curl --location $server/v1/medpipe/$processId/status"
    statusexec=`$getStatusUrl`
done
echo "The microservice is done. Results:"

getUrl="curl --location $server/v1/medpipe/$processId/predictions 2>/dev/null"
```

```
Result=$(eval "$getUrl")
echo "MED stats:"
echo $Result

getUrl="curl --location $server/v1/medpipe/$processId/tmh 2>/dev/null"
Result=$(eval "$getUrl")
echo "TMH:"
echo $Result

getUrl="curl --location $server/v1/medpipe/$processId/signal 2>/dev/null"
Result=$(eval "$getUrl")
echo "SIGNAL:"
echo $Result
```

I added a test file comprising two secreted proteins called “target.fasta”. However, feel free to try your proteins.

```
>Cp1002_0126a
MHFKTRMSLFCTATTAATSLAVASLQPAAAVEQPSNTIVSTIMLPTKATVTKTFTVSSTK
GTARADYSSNSITVQPGDTISVKIHSQGGYTEFSELTEFVPSVGRLHTESITFKEGDSGP
HPLKVAGWNATSQADRVTFRTNDGKPKAITLDTTLEYTYTVGVRATGDPSTRFQLSSSDS
NTVFTSASGPKIHVKKTLPSWLSGAFPGAIFDSLTNLLSPILRALNIL
>Cp1002_1802
MLFPSRFQGTFLKPLITAALAVFCVGFTPATAQVIPYTDPDGFYTSIPSAENTTPGTVLS
QRDVPMPVLDVLVKMKRIAYTSTHPNGFSTPVTGAVLLPTAPWRGPGPRPVALLAPGTQG
AGDSCAPSKLLTMGGEYEMFSAAALLNRGWTVAVTDYQGLGTPGNHTYMNRKAQGAALLD
LGRAITTLNLPDVNNHTPIIPWGYSQGGGASAAAAEMHRAYAPDVNVVLAYAGGVPANLL
SVSSSLEGTALTGALGYVITGMYEIYPEIREPIHNFLNTRGQVWLDQTSRDCLPESLLTM
PLPDTSILTVSGQRLTSLISDDVFQRAISEQQIGLTAPDIPVFVAQGLNDGIIPAEQARI
MVNGWLSQGADVTYWEDPSPALDKLSGHIHVLASSFLPAVEWAEQRLAALGQPTP
```

It’s important to note that complex names should be avoided. Otherwise, third-party software may trigger execution failures. As a precaution, I use the valifasta software to ‘clean’ protein files before running Medpipe. Please be aware that this important preprocessing step still needs to be implemented in the Medpipe microservice.

#### Medpipe microservice Overview

The source code for the microservice can be found on GitHub, and the project has received the name Medpipe microservice referencing the Medpipe tool that the microservice runs. The Medpipe microservice has three essential endpoints. The Medpipe script execution endpoint is used for directory preparation, temporary execution of files generated by Medpipe, and asynchronous execution of the script. Alternatively, it even makes use of H2 to record the status of the process and the generated directory for later consultation. The process status search endpoint is used to provide information about the status of Medpipe processing.

This functionality is used to find out if the processing of the Medpipe script has finished and if there were execution errors. The Medpipe prediction search endpoint is used to extract the results generated by Medpipe and return them to the user.

##### Execution of Medpipe

The Medpipe execution endpoint is triggered by an HTTP request using the POST verb in the /v1/medpipe/run route. It is an essential component of Medpipe microservice and is responsible for processing requests for the execution of the Medpipe script. It plays a central role in the automation of Medpipe processing, enabling users to submit files and parameters relevant to the execution of the Medpipe script.

##### Medpipe microservice technical details

Request type: POST Rota: /v1/medpipe/run Input Parameters:

- file: This parameter represents the Medpipe processing file. It is of the MultipartFile type, which is commonly used to handle file sending in web applications.
- cellWall: Measurement of cell wall thickness in amino acids.
- organism group: Group of organisms, currently accepted values are 0 for gram-negative or 1 for gram-positive bacteria.
- epitopeLength: Epitope length.
- E-mail: E-mail to send the results generated by Medpipe.
- membraneCytoplasm: A value of 1 considers predicting membrane and cytoplasmic proteins in addition to the default exported proteins. A zero value focuses only on the default.
- Directory Creation: First, use the buildDirectory function to build the Full path to a temporary directory where the results of the processing will be stored. This directory is generated in the folder "temp" with the name "MsMedpipe" concatenated with the timestamp to prevent duplication of Names.
- File Saving: Use the saveFile function to save the file received in the HTTP requests in the specified directory. That is essential because the Medpipe script requires access to the file.
- Process Control: In addition, a MedpipeControl object is created to track the process's status. This object is saved in the database and serves as a Medpipe processing progress monitoring tool.
- Medpipe script execution: We construct a shell command incorporating all the given parameters, including the file path, details on the cell wall,

group of organisms, and others. This command is executed through the Runtime.getRuntime().exec() function, which performs the Medpipe script in an asynchronous process.

- Status Update: The status of the process is updated based on the result of the script execution. If the run is successful, the status is set to done. If failure, the status indicates an error. It is worth noting that the runFileProcess function is annotated with @Async, which means that it runs on a separate thread. We accomplish that to ensure that the loop command does not block the main thread of the service and allows other requests to be processed efficiently, improving the scalability of Medpipe microservice. Code 1 is responsible for running Medpipe.

```
@Async
fun runScript(
    fileResult: String,
    cellWall: String,
    organismGroup: String,
    epitopeLength: String,
    email: String,
    membraneCitoplasm: String,
    medpipeControl: MedpipeControl
) {
    try {
        val command = "sh medpipe $fileResult $cellWall $organismGroup $epitopeLength $email $membraneCitoplasm"
        log.info("[ runScript ] - Start exec: ${LocalDateTime.now()} command: $command")
        val process = Runtime.getRuntime().exec(command)
        val processEnd = process.waitFor()
        val result = BufferedReader(InputStreamReader(process.inputStream)).readText()
        log.info("[ runScript ] - Time: ${LocalDateTime.now()} Result: $result")
        log.info("[ runScript ] End process: $processEnd")
        unzipFileResult(medpipeControl.directory)
        updateStatus(Status.FINISHED, medpipeControl)
    } catch (e: Exception) {
        updateStatus(Status.ERROR, medpipeControl)
        throw RuntimeException(e.message)
    }
}
```

Figure 1: Code 1

##### Search Processing Status

Status lookup is an endpoint that handles HTTP GET requests on the route /v1/medpipe/{id}/status. Its main purpose is to check the status of a Medpipe process based on the ID provided as a parameter in the Uniform Resource Locator (URL) of the request. Its operation is relatively simple.

Request Type: GET Rota: /v1/medpipe/{id}/status Input Parameter

- id: Parameter related to the processing of Medpipe, with which it is possible to search in the base H2 status information and file generation directory.

- Receiving the ID: The function receives the process ID as part of the URL of the HTTP request. The Spring Framework automatically extracts this ID due to annotation @PathVariable. - Status Query: The function invokes the findStatusProcess method of the MedpipeService service, passing the ID as an argument. This call to service is responsible for querying the status of the process in the database.
- Result Return: The status of the process, which is returned by the service, is then returned because of the getStatus function. The status is of type Long, allowing it to return an integer value representing the status of the process. This status can assume three pre-established values.
- Zero: Indicates that the process has been completed successfully.
- 1: Indicates that the process is still running and should wait.
- -500: Indicates that there was an error in the processing.
- -404: Indicates that no one was found in the database process referring to the ID received by the requisition. The endpoint gives users the ability to access information in real-time on the progress of Medpipe processes. It plays a key role in the transparency and monitoring of Medpipe script data processing.

Code 2 is responsible for fetching the processing status.

```
fun findStatusProcess(id: Long): Long? {
    log.info("[ findStatusProcess ] - ID: $id")
    val result = medpipeControlRepository.findById(id)
    return if (result.isPresent) {
        result.get().status?.statusCode
    } else {
        Status.NOT_FOUND?.statusCode
    }
}
```

Figure 2: Code 2

##### Searching for Medpipe predictions

The fetching of forecasts generated by Medpipe is an endpoint that handles HTTP requests using the GET verb in the /v1/medpipe/{id}/predictions route. The goal is to allow customers to view the predictions generated by Medpipe.

Request Type: GET Rota: /v1/medpipe/{id}/predictions Input Parameter

- id: Parameter related to the processing of Medpipe, allowing searching in the base H2 status information and file generation directory.
- Receiving the ID: The function receives the process ID as part of the HTTP request's URL. The Spring Framework automatically extracts this ID due to the @PathVariable annotation. The received ID is used in the

getPrediction function to get the processing predictions from the Medpipe script based on the ID provided.

- Process fetching: Internally, the getPrediction function uses the findProcess function to retrieve a MedpipeControl object based on the id. If the process is not encountered, an exception is thrown, indicating an HTTP 404 (NOT FOUND) status.
- File Search: After searching in the database, the MedpipeControl object contains the store directory with predictions.
- Result assembly: With the file path obtained, the function reads the file of results line by line using a BufferedReader. Each line is processed and removed, including information such as gene ID, numerical prediction, and type of protein, and added to a StringBuilder instance. The getPredictions endpoint provides customers with the ability to access and download Medpipe processing results efficiently. It plays a crucial role in making the results available for subsequent analysis or local storage, making the Medpipe microservice a versatile and useful tool for users. Code 3 is responsible for fetching the predictions generated by Medpipe.

```
fun getPrediction(id: Long): StringBuilder {
    log.info("[ getPrediction ] - Start exec id: $id")
    val medpipeControl = findProcess(id) ?: throw RuntimeException(HttpStatus.NOT_FOUND.toString())
    val fileResult = medpipeControl.directory + "/" + ResultFile.TARGET_FASTA_RESULT_SORT.description
    log.info("[ getPrediction ] - File Result: $fileResult")
    var line = ""
    val result = StringBuilder()
    if (!FileUtil.fileExists(fileResult)) {
        return result
    }
    BufferedReader(FileReader(fileResult)).use { br ->
        while (br.readLine()?.also { line = it } != null) {
            val values = line.split(" ").toTypedArray()
            result.append(values[0] + " " + values[4].substring(4) + " " + values[6] + "\n")
        }
    }
    return result
}
```

Figure 3: Code 3

#### Medpipe microservice Installation

In this section, we detail the process of generating and installing the Medpipe microservice executable. The source code for the microservice can be found on GitHub.

##### Dependencies

- Java Virtual Machine: The Kotlin language's build process is in the Java Virtual Machine (JVM). The required version is 11, and the "JAVA HOME" environment variable must be configured.

- Maven: Maven, a build tool, needs to be properly configured to run. It is necessary to download version 3.8 or later. Configure the environment variable "M2 HOME" with the directory where the installation of the maven is, for example: "M2 HOME=/home/apache-maven-3.8". Then, include the "M2 HOME" variable in the System Path pointing to the maven bin folder, for example: "PATH=$PATH:$M 2 HOME/bin".

##### Compilation process

To compile the project, one needs to use the maven "mvn" command. Using the system command line, navigate to the root directory from the Medpipe microservice project and type "mvn clean install." This command will download the project's dependencies, compile its classes, and create a target folder, and inside it, we will have a generated ".jar" file. The generated file follows the terminology that was configured in the pom.xml of the project (Code 4). The result is the combination of the name and version fields generating the file "ms-medpipe-1.0.0-Release.jar".

```
<groupId>com.tcc</groupId>
<artifactId>ms-medpipe</artifactId>
<version>1.0.0-Release</version>
<name>ms-medpipe</name>
```

Figure 4: Code 4

##### Installation

After compiling the project, the generated file "ms-medpipe-1.0.0-Release.jar" should be copied to Medpipe's root folder. The command executes the project: "java-en ms-medpipe-1.0.0-Release.jar."

##### Pannotator Integration

In this section, we will describe how the Pannotator integrates with the Medpipe microservice using the endpoints. Pannotator and Medpipe are basically bash scripts that can be run from the command line or through their web interfaces. With the development of Medpipe microservice, Pannotator now runs Medpipe through an API call exposed on the internet. Figure 1S shows this integration. The execution of Medpipe is done by the /v1/medpipe/run route and has been included in the Pannotator after updating the target fasta file. The FASTA format is used to store nucleotide sequences and proteins.

Code 5 defines the medpipePostURL variable to store a curl command used to send a POST request to the /v1/medpipe/run route. To proceed with that, one needs to specify the URL, the FASTA file to be sent, and the parameters required

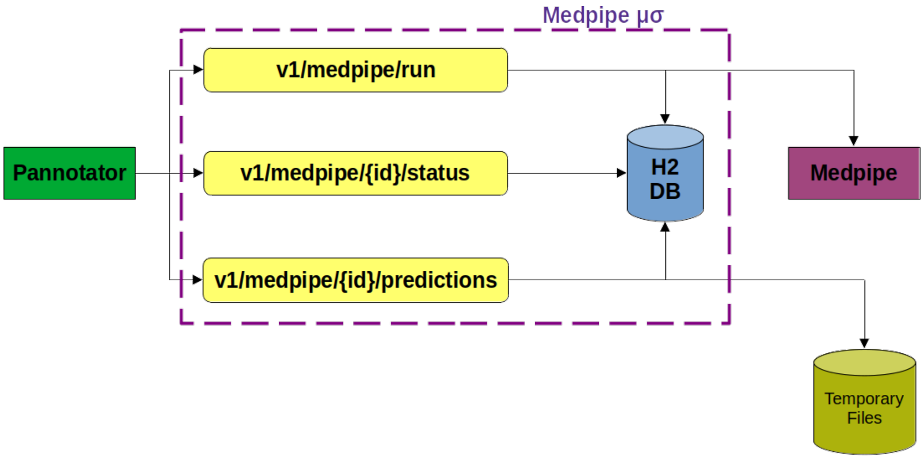


Figure 5: Figure 1S – Integration between Pannotator and Medpipe.

for the analysis. The command specified in the medpipePostURL variable is executed and stores the output in the medpipeexec variable. The API return is composed of Medpipe's processing ID. The ID returned by the execution service is used to obtain the information from the Medpipe process that has been initiated. The status endpoint receives this ID as a parameter and returns the status that is being processed.

```
medpipePostURL="curl --location localhost:8190/v1/medpipe/run \
--form 'file=@$dirfastaFile' \
--form 'cellWall=$cellWall' \
--form 'organismGroup=$organismGroup' \
--form 'email=$7'"
processId='$medpipePostURL'
```

Figure 6: Code 5

Code 6 executes the GET command to the Medpipe Processing Status API. After the Medpipe run is finished and the status is zero, the next step is to fetch the predictions generated by Medpipe.

```
# Pretty print getStatusUrl
echo "getStatusUrl ="
echo "curl --location"
echo "localhost:8190/v1/medpipe/$processId/status"

# Pretty print statusExec
echo "statusExec ="
echo "$getStatusUrl"
```

Figure 7: Code 6

Code 7 runs the fetching API for the projections generated by Medpipe. The Pannotator uses these predictions to enrich the execution of the results.

```
# Pretty print getPredictionsUrl
echo "getPredictionsUrl ="
echo "curl --location"
echo "localhost:8190/v1/medpipe/$processId/predictions"

# Pretty print medpipePredictions
echo "medpipePredictions ="
echo "$getPredictionsUrl"
```

Figure 8: Code 7

##### Documentation

To make the microservice easier to understand and use, it is essential to provide clear documentation. The following are details on how to interact with the endpoints of Medpipe microservice.

**Endpoint Documentation** Documentation for REST endpoints is available in the Swagger API, accessible under "http://localhost:[port configured]/swagger-ui.html". The documentation provides detailed information about each endpoint, as shown in Figure 2S, including the HTTP methods allowed and the expected responses.

**Logs** Ensuring proper visibility into microservice operations and performance is critical. To this end, logging resources were implemented for data analysis. Medpipe microservice generates detailed logs, recording relevant information about each operation, including data about incoming requests, responses sent, events, and any critical activities; an example can be found in Code 8.

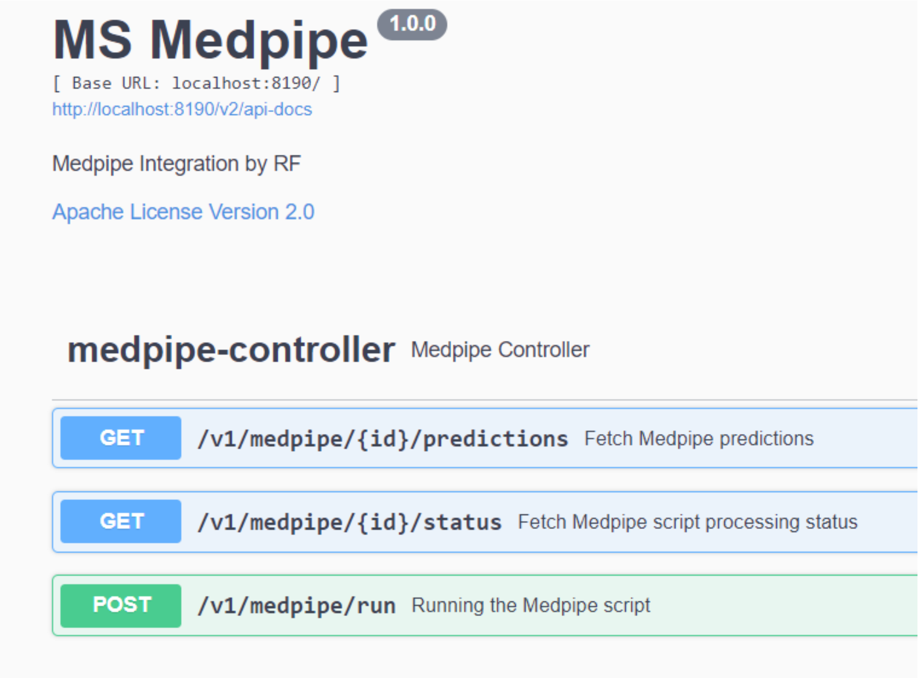


Figure 9: Figure 2S – Swagger API for Medpipe $\mu$ $\sigma$ documentation.

```
# [ runFileProcess ] - Init run ...
# [ runFileProcess ] - directoryRoot : /tmp/DB1703543511
# [ saveFile ] - Start - File : target.fasta directoryRoot: /tmp/DB1703543511
# [ saveFile ] - Result File : /tmp/DB1703543511/target.fasta
# [ runFileProcess ] - fileResult : /tmp/DB1703543511/target.fasta
# [ saveControl ] - Start - medpipe process: DB1703543511
# [ runFileProcess ] - process: com.tcc.medpipe.MedpipeControl@22671154
# [ runFileProcess ] - script terminated: /tmp/DB1703543511;1
# [ runScript ] - Start exec: 2023-12-25T19:35:20.242966 command: sh medpipe /tmp/DB1703543511/target.fasta 50 1 9
# [ runScript ] - Time: 2023-12-25T21:19:27.948890 Result: 2) Cell wall thickness for local subcellular prediction of /tmp/DB1703543511/target.fasta
# 3) Bacterial Gram for local subcellular prediction of /tmp/DB1703543511/target.fasta
# 4) Epitope length for MHC prediction of /tmp/DB1703543511/target.fasta
# 5) Eliminate extra text from fasta headers /usr/local/medpipe
# 6) GRAMP 1
# 7) Prepare to run surfG. Configuring current directory of installation
# MEDPIPE results:
# mygenome_00779 n =84
# [50 - avg ( aff )]=32.67 d =131 MED =20.94 FOLD =1.56 PSE
```

Figure 10: Code 8
